# Supplemental water modulates pollinator visitation on resource-rich plant species

**DOI:** 10.64898/2026.10.11.757154

**Authors:** Jaxi E. Cohen, Skye Austin, Angelica M. Sauceda, Rachel M. Davis, Emily B. Griswold, Rachel L. Vannette, Lauren Azevedo-Schmidt, Emily K. Meineke

**Affiliations:** Department of Entomology and Nematology, University of California, Davis, California, USA; Department of Evolution and Ecology, University of California, Davis, California, USA; UC Davis Arboretum and Public Garden, University of California, Davis, California, USA

**Keywords:** bumblebees, climate adaptation, drought, floral abundance, irrigation timing, native bees, urban gardens, water conservation

## Abstract

Urban green spaces provide important floral resources for pollinators, and supplemental watering can improve the abundance and quality of these resources. However, increasing frequency and intensity of drought under climate change create tradeoffs between ecological benefits and water conservation. Manipulating the timing and duration of irrigation may offer ways of enhancing floral traits that support pollinators while moderating water use. Focusing on two resource-rich native plant species, *Phacelia californica* and *Helenium puberulum*, we examined how the addition of supplemental water in spring and summer affects floral traits and pollinator visitation. We examined mechanisms that may govern the effects of supplemental watering on pollinator visitation, namely the number of flowers, nectar volume, nectar sucrose concentration, and corolla length. Plant individuals receiving more water were visited by more pollinators. In both plant species, this was apparently mediated by floral abundance; individual plants with more flowers received more visitors, though effects of watering varied slightly between plant species and insect visitor taxa. Water additions in spring increased pollinator visitation per plant on *P. californica*, but visitation rates per flower did not differ among treatments. Responses among pollinator groups varied, with effects driven primarily by sweat bees (family: Halictidae) and bumblebees (*Bombus* species), while honeybee visitation was consistent across treatments. Our results suggest that insects prefer to visit supplementally watered *P. californica* plants. Because honeybee visitation did not vary by treatment, supplemental water may disproportionately benefit native pollinators, including bumblebees, which are imperiled. Similar to *P. californica, H. puberulum* plants watered in spring and summer had the most flowers and the most floral visitors per plant and per flower. Spring-watering *H. puberulum* did not have a significant effect on flower production but did increase insect visitation per flower. Overall, our results emphasize that irrigation and other avenues of water addition are key drivers of flower production and insect visitation, such that homeowners who wish to conserve water must weigh effects on pollinators with the need to conserve water in arid climates.

## Introduction

Urban green spaces are important refuges for pollinators that can provide diverse floral resources (Baldock et al., 2019). Smaller green areas can also act as stepping stones that enable pollinator movement between larger areas (Dearborn & Kark, 2010). As agricultural expansion and intensification reduce floral resources in surrounding landscapes (Ollerton et al., 2014; Scheper et al., 2014), urban green areas can provide important forage and nesting habitats for pollinators (Frankie et al., 2009; Gunnarsson & Federsel, 2014). However, green spaces support pollinators to variable extents, and qualitative differences between urban habitats can modulate the number and diversity of pollinators (Baldock et al., 2019).

Urban refuges can be managed to increase pollinator abundance through diverse mechanisms such as plant species selection, sun exposure, and irrigation (Majewska & Altizer, 2018). As the global climate continues to change, these variables provide opportunities in which stakeholders, including homeowners, can adapt management to support urban pollinators. In particular, the provision of supplemental water through irrigation is often crucial to plant survival and growth. Water availability influences flowering time, which can affect the extent of overlap between flowering and peak pollinator activity (Neil et al., 2014; Kuppler & Kotowska, 2021). Supplemental watering can increase floral abundance and size, as well as affect the volume and concentration of nectar (Descamps et al., 2018). Such effects of water availability can mediate the signaling and nutritional value of floral rewards, leading to changes in pollinator visitation (Carroll et al., 2001; Descamps et al., 2021).

As climate change intensifies the frequency and duration of drought (Diffenbaugh et al., 2015; Pachauri et al., 2014) and reduces humidity during dry seasons (McKinnon et al., 2021), the capacity of urban habitats to support pollinators may increasingly depend on how changes in water availability affect the plant traits that mediate pollinator interactions. Urban irrigation practices vary widely, but even when managed water resources are limited, green spaces frequently rely on irrigation to support environmental benefits, such as floral resources for pollinators (Fam et al., 2008). Maintaining these benefits can come at the cost of increased water use, and the irrigation of green spaces represents a major component of household water usage in arid regions (Mini et al., 2014; Saher et al., 2023). At the same time, many insect pollinator populations are declining (Goulson et al., 2015), and decisions about urban irrigation must account for trade-offs between ecological benefits and water conservation efforts.

Variation in the timing and duration of irrigation is a potential pathway to manipulate floral traits to serve pollinator communities while still minimizing water use. Water availability during flowering can have stronger effects on floral traits such as nectar volume, floral abundance, and corolla size than conditions before flowering (Kuppler & Kotowska, 2021). Temporal variation in irrigation may also decouple flowering periods from pollinator activity, as different stages in plant development may respond distinctly to changes in water availability (reviewed by Descamps et al. 2021; Omidi, 2009). Utilizing these differences may allow management strategies to provide supplemental water only when most beneficial.

In California, adaptations to a Mediterranean climate allow many native plant species to cope with periods of seasonal drought, including maintaining floral rewards during water stress (Nardini et al., 2014). From an urban management perspective, this means that native species might not require supplemental watering in every season to maintain floral resources. Selecting these species can lower the need for supplemental watering (Belaire et al., 2023), while still maintaining resources for pollinators (Williams et al., 2015). Understanding how temporally distinct watering treatments influence drought-adapted plants is essential for supporting pollinator communities under future climate and water management scenarios.

Here, we quantified the effects of supplemental water on two native plant species that support diverse and abundant pollinator communities in urban areas in California. We established a novel experimental framework called the Climate Adaptation Living Lab (CALL) garden at the UC Davis Arboretum and Public Gardens. We manipulated the timing of water availability and the seasons in which plants were watered via drip irrigation, similar to that which is installed by regional homeowners. Treatments differed by irrigation timing and duration: (1) watered during establishment only, (2) watered during spring, (3) watered during spring and summer, and (4) well-watered throughout. We tested two non-exclusive hypotheses: (1) Increased Floral Resources Hypothesis: Watering increases the number of flowers, leading to increased pollinator visitation, (2) Nectar Dilution Hypothesis: Supplementally watered plants produce nectar with lower sucrose concentration, which reduces pollinator visitation. Due to the water treatments’ variation in timing, our predictions under these hypotheses differed between the spring and summer sampling periods.

## Materials and methods

### Focal species

We studied two California native perennial herbs adapted to Mediterranean seasonal drought: rock phacelia *Phacelia californica*, a spring-flowering plant chaparral and coastal scrub with bell-shaped flowers borne in cymes, and sneezeweed *Helenium puberulum*, a summer-flowering plant of seasonally moist meadows and riparian wetlands with flower heads of hundreds of florets (Calflora, 2026; Natural Resources Conservation Service, 2026). *Phacelia* species are nectar- and pollen-rich and frequently visited by bumblebees (*Bombus spp.*), honeybees (*Apis mellifera*), and solitary native bees (Bruckman & Campbell, 2014). Both species are commonly recommended for pollinator gardens (Pawelek et al., 2015).

### Experimental design

The study was conducted at the Climate Adaptation Living Lab (hereafter, the “CALL Garden”) located in the UC Davis Arboretum and Public Gardens. We established 40 1.52 x 1.52 m plots arranged in 10 rows of four, built as 15.24 cm deep wooden frames filled with field soil (Yolo silt loam) and topped with fir mulch, similar to regional home garden beds. Plants were transplanted into plots on December 4, 2024. Each plot contained one individual of five native species: *Phacelia californica, Helenium puberulum*, *Artemisia douglasiana, Salvia mellifera,* and *Symphoricarpos albus* var. *laevigatus*. Plots were irrigated with surface drip lines (emitters every 30.48 cm, 2.27 L h^-1^) controlled by treatment specific schedules (Supplementary Methods).

Plots were randomly assigned to one of four water treatments (10 plots each): well-watered throughout, watered in spring, watered in spring and summer, watered in establishment only (hereafter “not watered”). All plots received uniform establishment watering beginning March 3, 2025, and treatment-specific watering began March 20, ending watering for the not-watered plots. Spring-watered plots stopped receiving water on June 20, 2025. All irrigated plots received water for 3 h every Monday and Friday (05:00 to 08:00). Plots were hand-weeded throughout.

The treatments active during each sampling period differed (Fig. 1). During *P. californica* sampling (April-May), the spring, spring-summer, and well-watered plots were all irrigated, so we compared watered plots (n = 30) with unwatered plots (n = 10). During *H. puberulum* sampling (June-July), we compared not watered plots (n = 10), spring-watered plots (n = 10, no longer irrigated), and spring-summer-watered plots (n = 20, spring-summer and well-watered plots, which received identical watering over this period). On July 16, an irrigation error watered all plots continuously, eliminating the treatment differences, so we excluded all observations after this date.

**Figure 1:**
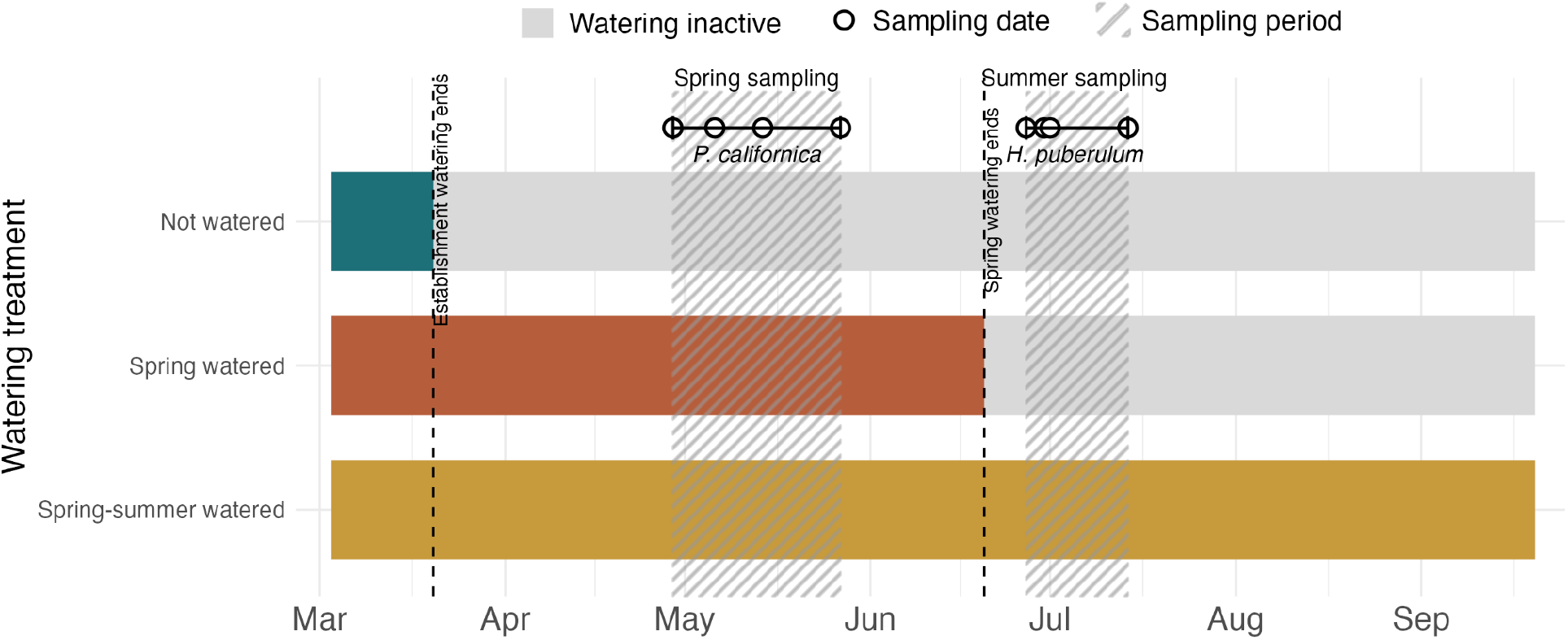
Timeline of watering treatments and sampling periods. Gray bars indicate when supplemental watering was inactive, while colored bars (corresponding to watering treatment) indicate periods when each treatment was receiving supplemental water. Vertical dashed lines indicate when changes in supplemental watering occurred. The diagonal striped regions indicate sampling periods, with specific sampling dates represented by circles. Well-watered throughout plots received the same water as spring-summer plots and are represented as one treatment.

### Visitation

#### Phacelia californica

We observed each plot for 3 min on April 29, May 6, May 14, and May 27, 2025, between 12:00 and 16:00 under sunny, low-wind conditions. Observations were conducted on the whole plant and classified visits to visually distinguishable taxonomic groups: *Bombus, Apis mellifera*, small, large, and metallic Halictidae*, Xylocopa*, Lepidoptera, and Diptera. Most visits were by *Bombus*, *A. mellifera*, and small Halictidae. A single visit was recorded when an insect made contact with the flower.

#### Helenium puberulum

We observed each plot for 3 min on June 27, June 30, July 1, and July 14, 2025, between 09:00 and 16:00 under low-wind conditions. Sampling was conducted in both morning and afternoon to capture the earlier shift in pollinator activity during summer. Due to the large number of flowers per plant, observations were conducted on 10 apical flower heads per plant. All visits were Halictidae and were recorded as a single group. A single visit was defined the same as with *P. californica*.

### Floral traits

In May 2025, we sampled nectar from five flowers per *P. californica* plant, taken from cymes bagged before opening to exclude pollinators. Collection occurred before 11:00 to limit evaporation. Nectar volume was measured with 10 µL microcapillary tubes and nectar concentration was measured using a handheld refractometer. Corolla length was measured under a dissecting microscope on three flowers per plant collected over 5 days within an 8-day period in May. Nectar and corolla traits were not measured for *H. puberulum* due to small size of flowers. We counted open flowering units per plot (inflorescences for *P. californica*, flower heads for *H. puberulum*) in approximately weekly phenology surveys from April 13, 2025 (Table S1).

### Sun exposure

Surrounding foliage shaded some plots in the morning. We classified each plot as sun or shade at 09:00, when pollinator activity begins, using a time-lapse recorded on June 26, 2025. Flower surface temperature measured with an infrared thermometer was higher in sun than in shade plots and did not differ among watering treatments (Table S4).

### Soil moisture

We measured soil water content and water potential at 30 cm depth (METER TEROS 11 and 21 sensors logged every 4 h; METER Group, Inc. Pullman, WA, USA) in two plots per treatment in May 2025. We summarized data as means ± SD per treatment before and after June 20, when spring watering ended.

### Statistical analyses

We analyzed each species separately in R version 4.5.2. (R Core Team, 2025). Observed visit counts were integers, so we modeled them with Poisson generalized linear mixed models (GLMMs; R package lme4; Bates et al., 2015). Responses derived from counts by division or multiplication are continuous, so we analysed them with linear mixed models (LMMs) with Gaussian errors (R package lme4; Bates et al., 2015). We considered structural equation modeling (Grace et al., 2015), but preferred mixed models to include random effects. Plot, which equals the individual plant, was specified as a random effect for repeated measures. The model structure for each response is listed in Table S4.

Response variables differed between species because of the sampling methods, so we calculated visitation at two scales for each. For *P. californica*, we analysed visits per plant by taxonomic group (*Bombus, A. mellifera*, small *Halictidae*), total visits, and taxonomic richness (number of groups observed) as Poisson GLMMs. Visits per flower were visits per plant divided by the number of flowers on the phenology survey closest in time, analyzed with Gaussian LMMs. We did not calculate richness per flower because richness is bounded and does not scale with flower number. For *H. puberulum*, we analysed visits per flower as Poisson GLMMs. Visits per plant were extrapolated as visits per flower multiplied by the number of flowers on the closest survey, analyzed with Gaussian LMMs.

For each response, we fit two kinds of models. Treatment models included watering treatment (watered vs. not watered for *P. californica*; not watered, spring-, and spring-summer-watered for *H. puberulum*), date, their interaction, morning sun exposure, and time of day. Mechanistic models included, for *P. californica*, number of flowers, mean corolla length (averaged across three flowers), sun exposure, time of day, and either mean nectar volume or sucrose concentration. Mean nectar volume and sucrose concentration were correlated (r = -0.53), and were included in separate models: the Nectar Volume (hereafter, “NV”), and the Nectar Sucrose Concentration (“NSC”) models. For *H. puberulum*, they included number of flowers, sun exposure, and time of day. Date was excluded from mechanistic models to avoid collinearity with flower number (*P. californica*: r = 0.54; *H. puberulum*: r = -0.44). Flower number and nectar traits were matched to the nearest visitation date. We tested the effects of watering on flower number, nectar volume, and sucrose concentration with LMMs (Table S4), and on flower surface temperature with an LMM including plot and date as random effects.

Pairwise contrasts among treatments were estimated with emmeans (Lenth, 2025), using Tukey adjustment for the three treatments of *H. puberulum*. *Phacelia californica* had two treatment levels, so a single unadjusted contrast was used. Contrasts from the log-link GLMMs are reported as rate ratios, and contrasts from the LMMs are reported as differences in visits per plant per 3 min. Coefficient *p*-values in Tables 1 and S3 are not adjusted for multiple comparisons.

### Permits and permissions

The study was conducted in a research garden managed by the authors’ institution, and no permits were required for non-invasive observation of insects.

## Results

### Effects of supplemental water on visitation

#### Phacelia californica

##### Treatment models

Spring watering increased total visitation per plant 1.40-fold (*β* = 0.34, *P* = 0.008; Table 1; Fig. 2), but did not change visitation per flower (*P* = 0.83), which declined over time and with time of day. Bumblebee and Halictidae visitation per plant were 1.48 times and 1.68 times higher on spring-watered plots (*β* = 0.39, *P* = 0.006; *β* = 0.52, *P* = 0.02), while honeybee visitation did not differ (*P* = 0.50; Fig. 4). No group differed per flower. The increase in Halictidae per-plant visitation over time was greater for spring-watered plants (*β* = 0.39, *P* = 0.004). Sun exposure and time of day did not affect any group. Taxonomic richness per plant was not affected by treatment or other predictors (*P* = 0.58).

**Table 1:**
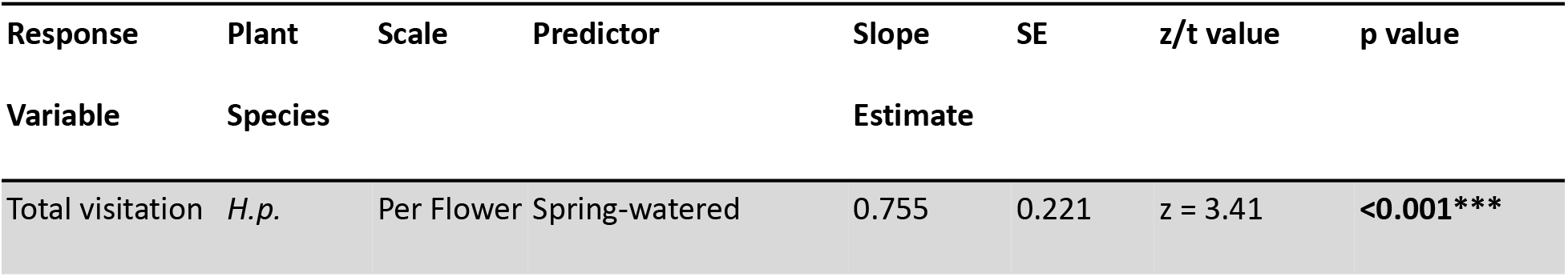
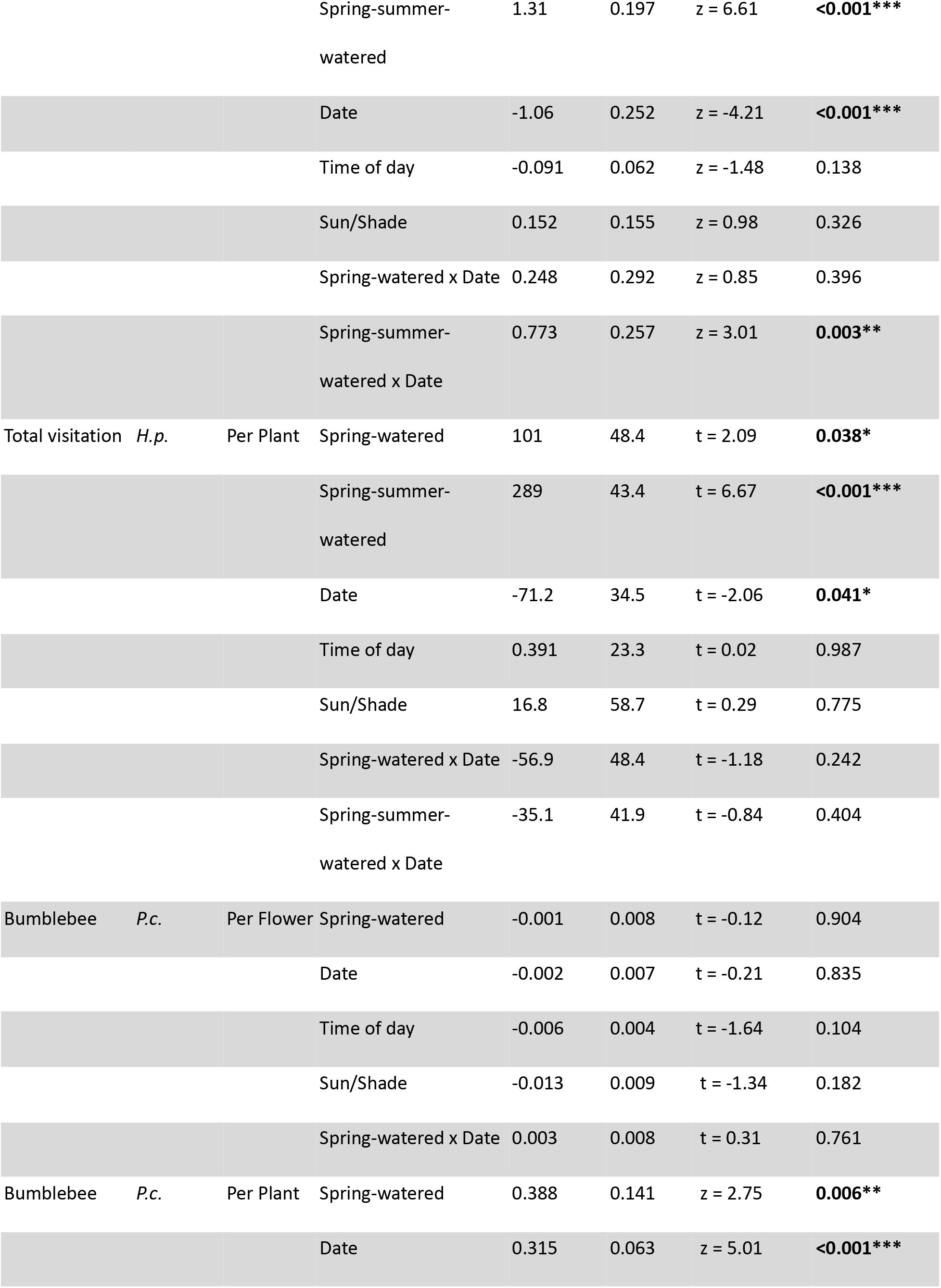
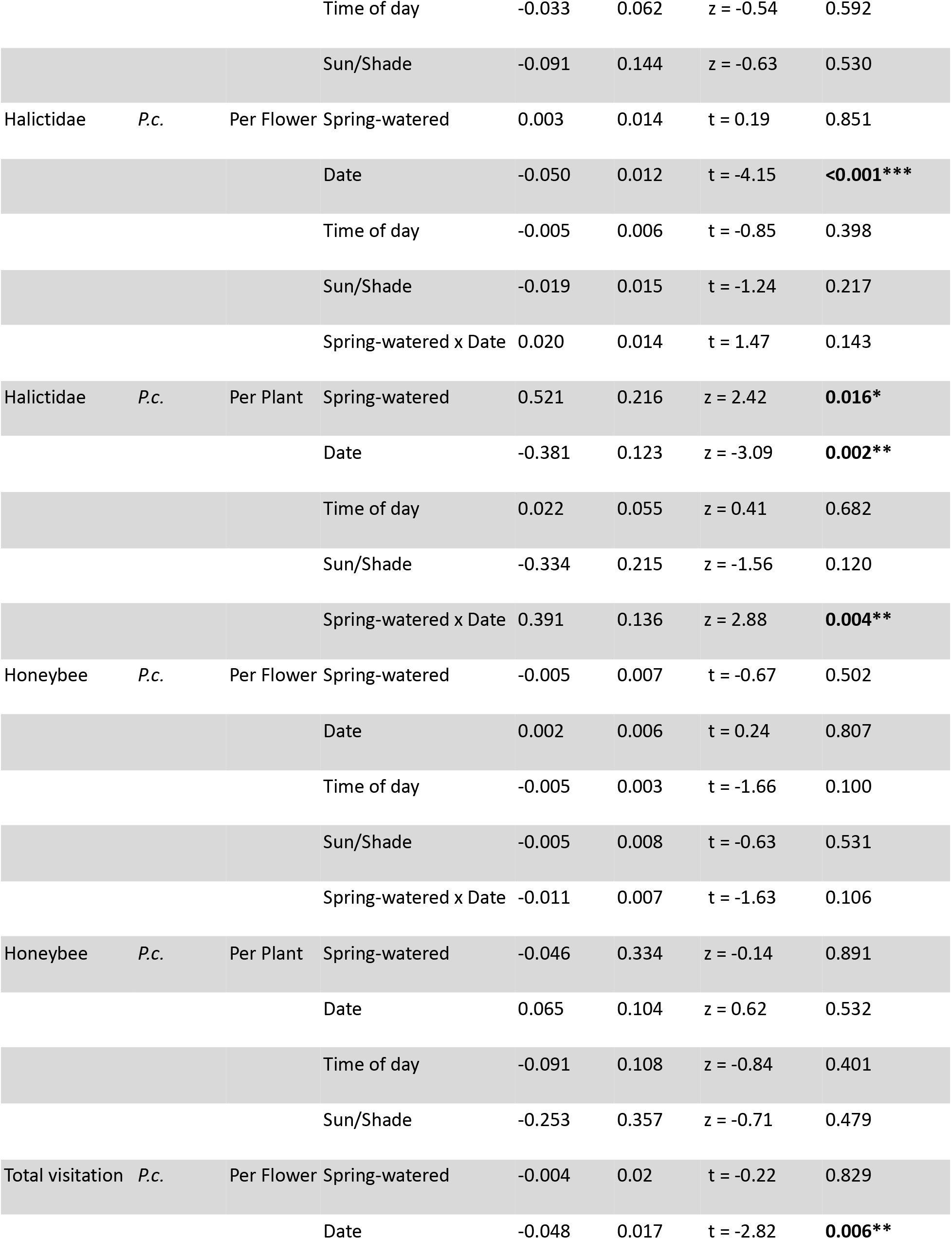
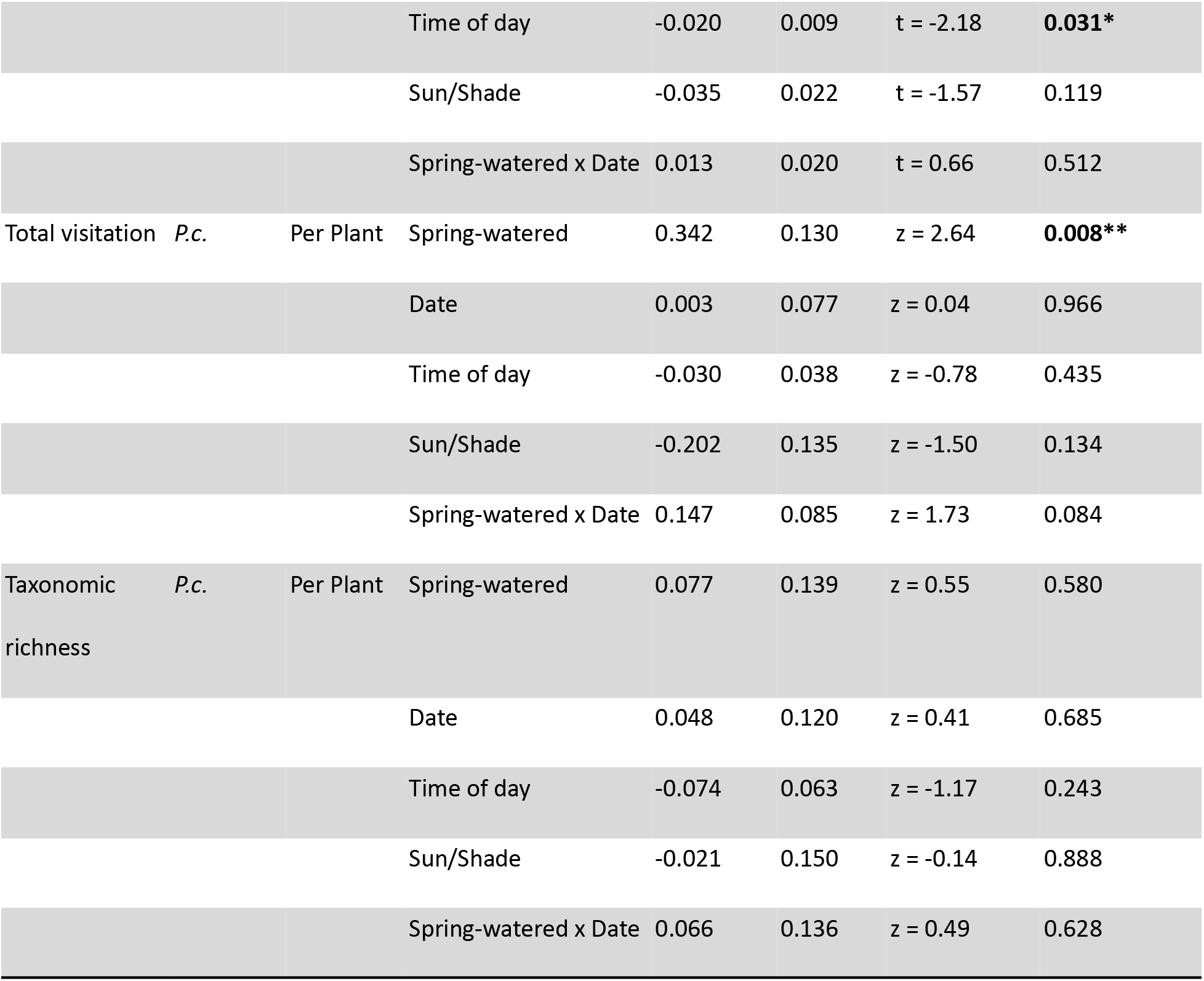
Treatment model results. *P. californica* per plant and *H. puberulum* per-flower visitation: Poisson GLMMs (log-scale estimates, *z*). *P. californica* per flower and *H. puberulum* per-plant visitation: Gaussian LMMs (estimates in visits per plant per 3 min, *t*). *P*-values are from model coefficients; Tukey-adjusted contrasts are in the text. * *p* < 0.05, ** *p* < 0.01, *** *p* < 0.001. *P.c., Phacelia californica; H.p., Helenium puberulum*.

**Figure 2:**
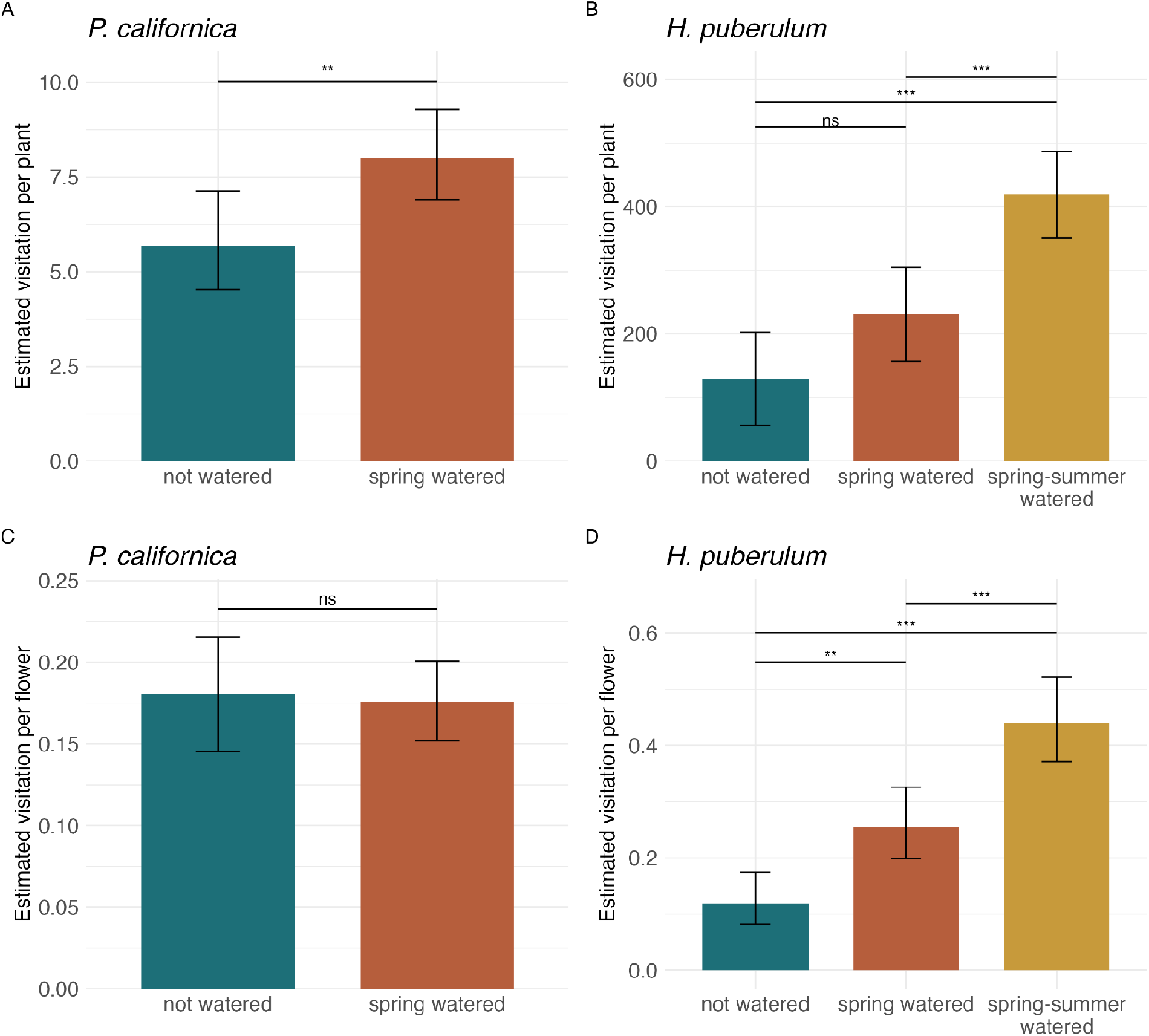
Estimated visitation per plant (a, b) and per flower (c, d) per 3-min observation period of *P. californica* (a, c) and *H.puberulum* (b, d). Bars are model-estimated marginal means with 95% CIs. Brackets show pairwise contrasts (Tukey-adjusted for *H. puberulum* (b, d): ns *p* > 0.05, * *p* < 0.05, ** *p* < 0.01, and *** *p* < 0.001.

##### Mechanistic models

Visitation per plant increased with the number of flowers for total visitation and every taxonomic group (all *P* < 0.03; Fig. 3, Table S3), except honeybees in the NV model (*P* = 0.07), as did taxonomic richness (NV: *β* = 0.17, *P* = 0.003; NSC: *β* = 0.17, *P* = 0.003). Morning sun exposure was associated with lower total visitation and, in the NV model, lower Halictidae visitation (*β* = -0.46, *P* = 0.04). Bumblebee visitation declined through the afternoon (NV: *β* = -0.13, *P* = 0.03; NSC: *β* = -0.12, *P* = 0.03). Honey bee visitation was lower on plants with higher nectar sucrose concentration (*β* = -0.35, *P* = 0.02). Longer corollas were marginally associated with higher total visitation (NSC: *P* = 0.06; NV: *P* = 0.05). Nectar volume did not affect visitation.

**Figure 3:**
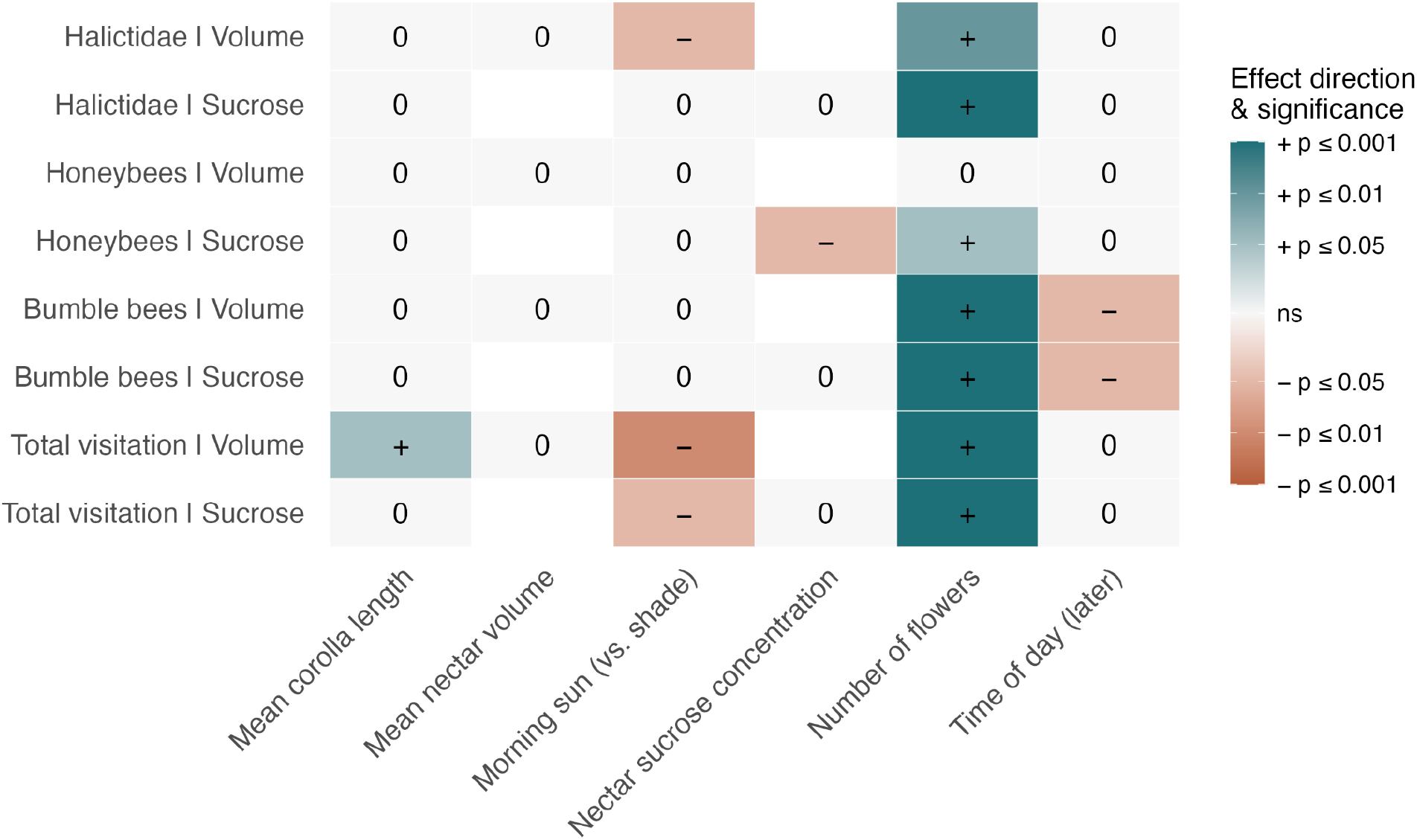
Effects of predictors on per-plant visitation to *P. californica* (Poisson GLMMs; Table S4). Each tile shows the direction and significance of a predictor in a Poisson GLMM of per-plant visitation (Table S3). Nectar volume and nectar sucrose concentration were included in separate models. Morning sun is exposure at 9:00 AM relative to shade.

**Figure 4:**
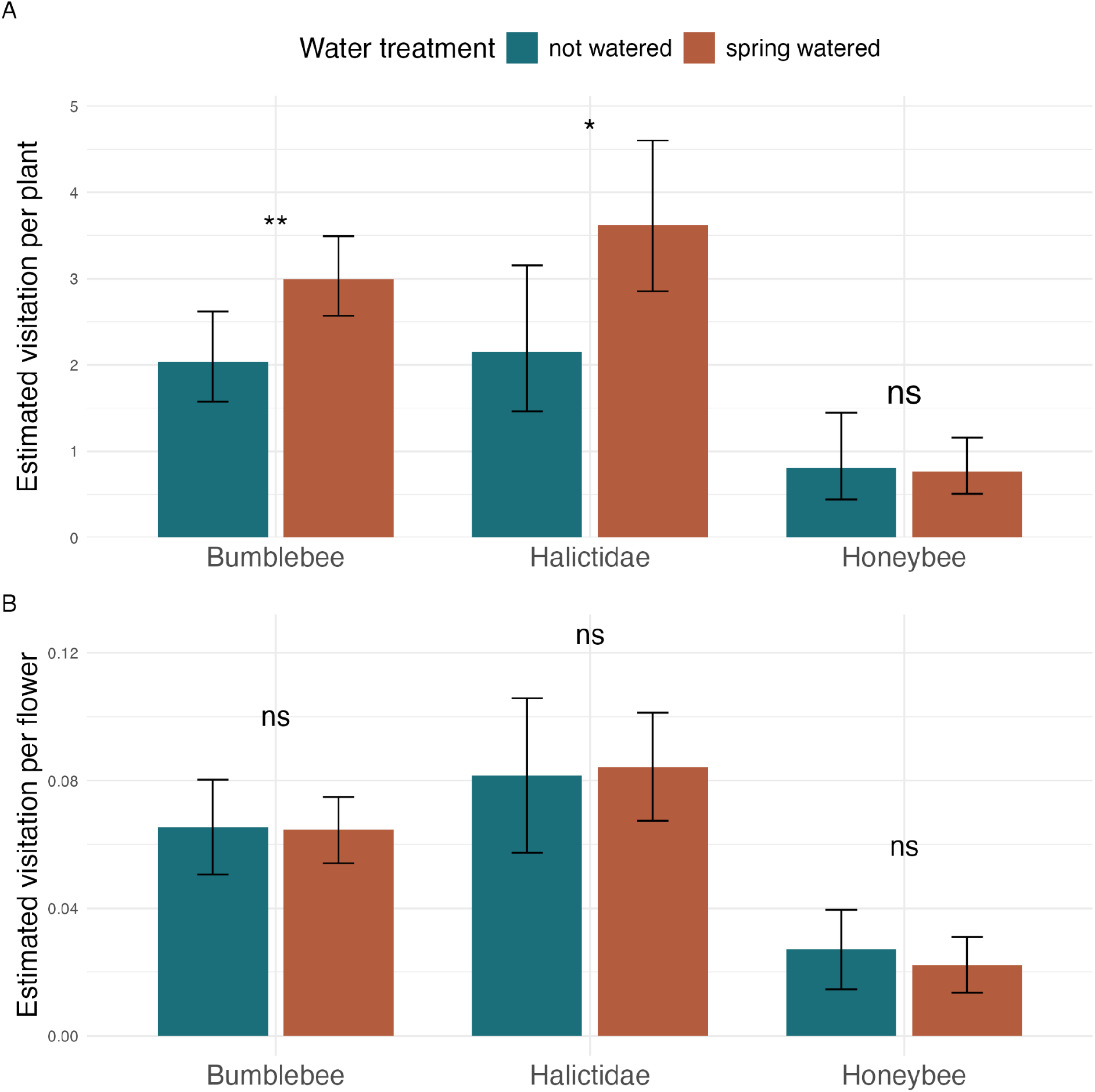
Estimated visitation by taxonomic groups to *P. californica* (a) per plant and (b) per flower per 3-min observation period. Bars are model-estimated means with 95% CIs. ns *p* > 0.05, * *p* < 0.05, and ** *p* < 0.01.

#### Helenium puberulum

##### Treatment models

Spring-summer-watered plots had significantly higher visitation at both the plant and flower level, while spring-watered plots had higher visitation only at the flower level. Total visitation per flower to spring-summer-watered plots was 3.69 times higher than to unwatered plots (*β* = 1.31, Tukey-adjusted *P* < 0.001; Table 1; Fig. 2), and 1.73 times more than spring-watered plots (Tukey-adjusted *P* < 0.001). Total visitation per plant was about 289 visits per plant per 3 min higher in spring-summer-watered plots than unwatered plots (Tukey-adjusted *P* < 0.001) and about 188 visits higher than in spring-watered plots (Tukey-adjusted *P* < 0.001). Total visitation per flower to spring-watered plants was 2.14 times higher than unwatered plots (*β* = 0.76, *P* < 0.001), while total visitation per plant was not significantly higher (Tukey-adjusted *P* = 0.11). Visitation to unwatered plots decreased later in the sampling period (Per flower: *β* = -1.06, *P* < 0.001; Per plant: *β* = -71.2 visits per SD of date, *P* = 0.04). The increase in total visitation per flower over time was greater in spring-summer-watered (*β* = 0.77, *P* = 0.003), but not visitation per plant, and spring watering did not alter it, indicating that the effects of supplemental watering became stronger at the flower level as summer progressed. Time of day and morning sun exposure had no effect.

##### Mechanistic models

Visitation per flower and per plant increased with more flowers (Per flower: *β* = 0.39, *P* < 0.001; Per plant: *β* = 141, *P* < 0.001; Table S3). Time of day and morning sun had no effect.

### Plot-level sun exposure

Plants exposed to sun in the morning had increased surface temperature of flowers compared to shaded plants (*β* = 8.02, *P* < 0.005). Water treatment did not affect flower surface temperature (*P* = 0.42).

### Effects of water treatments on floral traits

#### Phacelia californica

Spring-watered plants produced more flowers than unwatered plants (*β* = 9.17, *P* = 0.04), with a steeper increase over time (*β* = 0.59, *P* < 0.001). Watering did not affect nectar volume or sucrose concentration. Nectar sucrose was higher in plots with morning sun exposure (*β* = 6.34, *P* = 0.04), and nectar volume and sucrose concentration were negatively related.

#### Helenium puberulum

The number of flowers increased over time (*β* = 10.7, *P* < 0.001). Spring-summer watering increased flower number (*β* = 64.4, *P* = 0.03), but not spring watering (*P* = 0.18), with steeper increases over time for both watered treatments (spring-summer: *β* = 2.98, *P* < 0.001; spring: *β* = 1.78, *P* = 0.05).

### Effects of water treatments on soil moisture

Soil was wetter in watered than unwatered plots (Table S2): before June 20, water potential averaged -445 kPa in watered plots versus -2021 kPa in unwatered plots, and after June 20, -50, -2631, and -3297 kPa in spring-summer, spring, and unwatered plots.

## Discussion

### Overview

This study aimed to determine how the modulation of plant water availability can affect the resources available to insects. Our results support the Increased Floral Resources Hypothesis; regardless of the plant species, individual plants that were watered more produced more flowers and received more floral visitors, though important nuances emerged among plant and insect taxa. On *P. californica*, bumblebees and Halictidae responded to water treatments while honeybees did not, indicating that water additions may have a stronger and more positive impact on taxa that we presume are native, a subset of which are imperiled. We did not find support for the Nectar Dilution Hypothesis as watering did not affect nectar sucrose concentrations.

### Effects of floral abundance and size

The scale at which pollinators responded differed between plant species. In *P. californica*, watering increased visitation only at the plant level, indicating that pollinators responded primarily to whole-plant signals, namely the total number of inflorescences per plant. Supplemental watering increased visitation at both the plant- and flower-levels in *H. puberulum*, suggesting that it may have affected both plant- and flower-level attractiveness or rewards.

Spring-watered *P. californica* plants received greater visitation at the plant level, but visitation per flower did not vary across watering treatments. This suggests that pollinators responded to changes in the detection of the whole plant, instead of individual flower attractiveness or rewards. Consistent with this finding, the increase in total visitation per plant of *P. californica* was driven by the number of flowers. Plants with more flowers can be easier to detect due to their larger size or visual distinctiveness (Ohashi & Yahara, 2001). As spring-watered plants had a greater number of flowers, our results support a model in which supplemental watering indirectly increased pollinator visitation through greater flower production.

Taxonomic richness did not differ among watering treatments but increased with the number of flowers, meaning supplemental watering did not change which visitor groups were present. Because richness was assessed using coarse, visually identified groups and richness generally increases with the number of visits, larger floral displays may have simply provided more opportunities to detect additional taxonomic groups.

Similar to *P. californica*, watering treatment affected total visitation at the plant level in *H. puberulum* and was also mediated by floral abundance. However, our interpretation of *H. puberulum* results must account for potential inaccuracy in plant-level visitation estimates, as they were calculated by multiplying flower-level data by the number of flowers, and visitation could not be monitored at the plant level (across all flowers) due to high floral abundance. Taking this into account, spring-summer watering, but not spring watering, increased total visitation per plant. Plants watered in spring and summer had significantly greater visitation per plant than both spring-watered and unwatered plants, suggesting pollinators preferred spring-summer-watered plants due changes in the attractiveness of the whole plant. This is further supported by the increase in floral abundance in spring-summer-watered plants, but not spring-watered plants, relative to unwatered plants. The effect of water treatment on total visitation per plant was driven by the number of flowers, such that spring-summer watering increased flower production, which then resulted in more visitation by pollinators.

Spring watering alone did not affect total visitation per plant of *H. puberulum*. Consistent with these results, spring-watered plants did not have a significantly greater number of flowers compared to unwatered plants. This suggests that watering during spring alone is not sufficient to maintain floral abundance or pollinator presence, and that continued watering throughout summer is crucial to maintain pollinator support. However, as described above, these results must be interpreted with caution due to our sampling methods.

The effect of watering treatment on visitation per flower varied by plant species. In *P. californica,* total visitation per flower did not increase with spring watering, and instead declined on plants with a greater number of flowers. This suggests that, despite plants with larger floral displays attracting more pollinators overall, visits were distributed among a greater number of flowers, reducing the probability of an individual flower receiving a visit. Per-flower visitation rate has been shown to decrease with the number of flowers for this reason in other studies (Akter et al., 2017; Klinkhamer & de Jong, 1989). Therefore, our results are consistent with a model in which spring watering increased the attractiveness of the plant as a foraging target without changing the attractiveness of individual flowers.

Unlike *P. californica*, both spring watering and spring-summer watering increased total visitation per flower on *H. puberulum*. This suggests that water additions increased the attractiveness and/or rewards of individual flowers. As we did not measure flower-level traits, we can only speculate as to what mechanism flower attractiveness changed by. Pollinators may prefer individual flowers based on traits such as nectar quality or floral scent, both of which can be affected by water availability (Kuppler & Kotowska, 2021). Spring-summer watering led to a greater increase in per-flower visitation than spring watering, relative to unwatered plants. This further supports our findings concerning per-plant visitation, highlighting the importance of watering during summer. While watering during spring only may increase visitation per flower, continued watering throughout summer provides an additional benefit. This is reinforced by our results concerning the interaction of watering treatment and date, as the effect of spring-summer watering on per flower total visitation was greater later in summer. There were no interaction effects of watering treatment and date for spring watering alone. As the summer progressed, and the longer spring-watered plants were without water, the greater the importance of watering during summer to pollinator presence.

Interestingly, per-flower visitation to *H. puberulum* was also driven by the number of flowers. Plants with greater floral abundance had greater visitation per flower, in contrast to visitation per flower of *P. californica*. This indicates that spring and spring-summer watering increased per-flower visitation to *H. puberulum* despite visits being distributed among a greater number of flowers. Our interpretation may be limited by our sampling methods, as we only observed visitation on the apical flowers of each plant. This may affect the relevance of floral abundance as a predictor of per-flower visitation, as inflorescence height relative to surrounding vegetation can affect pollinator detection (Gumbert & Kunze, 1999). Apical flowers may be more visible or accessible to pollinators due to greater vertical position, leading to increased pollinator visitation (Klečka et al., 2018). Therefore, the increase in per-flower visitation may not be representative of the greater number of flowers to distribute visits to. Further complicating our interpretation is our finding that floral abundance was not significantly greater in spring-watered plants, meaning that pollinators did not display a preference for spring-watered flowers due to plant-level floral abundance. Together, these results support the hypothesis that supplemental watering affected visitation of *H. puberulum* by both plant and flower-level mechanisms, leading to plant-level increases in spring-summer-watered plants only, and flower-level increases in both spring and spring-summer-watered plants.

In *P. californica*, in addition to the number of flowers, we found support that the effect of spring watering on total visitation may have been mediated by flower size. Plants with flowers that had longer corollas received greater total visitation per plant. Larger mean corolla length may increase the overall attractiveness of a plant if pollinators preferentially select plants with larger flowers. However, mean corolla length may also be associated with unmeasured characteristics such as plant size (Gómez et al., 2009), which can influence pollinator visitation (Klinkhamer et al., 1989). Future research could help determine whether the positive relationship between mean corolla length and visitation can be attributed to larger individual flowers or correlated plant characteristics.

### Effects of sun exposure and time of day

After noticing the variation in shade conditions across plots, we quantified morning sun exposure to determine if it played a role in the variation of plant traits and pollinator visitation. Morning sun exposure was associated with reduced total visitation per plant in *P. californica*. Sun exposure affects solar radiation and flower temperature, which can both directly alter floral resources such as nectar (Kilkenny & Galloway, 2008; Petanidou & Smets, 1996). Greater solar radiation and temperature can increase nectar sucrose concentration and volume due to greater evapotranspiration and photosynthetic activity, leading to changes in pollinator response (Plos et al., 2023). This finding may indicate that morning sun-exposed plants were less attractive to pollinators due to a change in plant traits. Equally likely is that the timing of pollinator activity was earlier in the plots exposed to sun earlier in the morning, and we missed the relevant visitation window, as our observation periods all took place from 12:00 to 16:00. Pollinators may have shown a preference for sun-exposed plants earlier in the day, and then moved on to forage from the remaining plants after depleting the preferred resources. Insect pollinators may forage sun-exposed plants first due to higher rates of nectar production (Petanidou, 2007), or the warmer microenvironment (Everaars et al., 2011; Herrera, 1995), as shade decreases temperature. Future studies could address whether variation in sunlight over the course of a day can increase the window of resource availability for pollinators.

Morning sun exposure did not affect total visitation of *H. puberulum* at the plant nor flower level. If the effect of sun exposure on visitation is mediated by nectar rewards, this difference between species may be attributable to the greater nectar-richness of plants in the genus *Phacelia* (Williams, 1996; Wróblewska, 2010). The effects of sun exposure may be dependent on both plant species and the floral trait mediating its effect. Additionally, our sampling period for *H. puberulum* included morning pollinator activity, therefore providing a more comprehensive view of pollinator activity that we did not capture in our sampling of *P. californica.* The lack of significant effect of sun exposure on floral visitors of *H. puberulum* may suggest that morning sun exposure may not affect overall visitation throughout the day, but instead the timing that plants are visited.

Time of day was particularly relevant for bumblebee visitors of *P. californica.* In addition to increasing with the number of flowers, bumblebee visitation per plant decreased over the course of the day (from 12:00 to 16:00). High temperatures in the late afternoon can negatively impact bumblebee foraging by decreasing foraging duration and the proportion of successful foraging bouts (Hemberger et al., 2022). Their large body size and adaptation to cooler climates make bumblebees vulnerable to high temperatures, as they struggle to shed the internal body heat necessary to maintain flight capacity (Martinet et al., 2015). Honeybee and Halictidae visitation did not vary by the time of day, further supporting that bumblebees are especially sensitive to temperature.

### Effects of supplemental water varies by insect taxa

While spring watering of *P. californica* increased total visitation per plant, this response was not consistent across taxonomic groups. The increase in visitation was driven primarily by bumblebees and bees in the family Halictidae, and, notably, honeybees displayed no preference for plants according to water input. However, the number of flowers increased visitation per plant across all three taxonomic groups. Honeybee visitation was therefore indirectly affected by watering treatment, as the number of flowers increased with spring watering.

Honeybee visitation may have shown looser ties to water treatment because of additional forces shaping honeybee foraging. Honeybees were the only taxonomic group affected by the nectar sucrose concentration of *P. californica*. Supplemental watering did not affect nectar sucrose concentration, which is consistent with previous studies (Descamps et al., 2018; Gallagher & Campbell, 2017) and indicates that variation in nectar quality contributed to honeybee foraging decisions independent of supplemental watering. Honeybees commonly use nectar sucrose concentration to distinguish between nectar rewards and alter their foraging preferences (Nicolson et al., 2013). Therefore, honeybee visitation was influenced by multiple characteristics of individual plants, while the relationship between supplemental watering and other taxa’s visitation may be more indicative of the treatment-level differences in floral abundance. This pattern suggests that the same environmental treatment can influence different pollinator taxa through distinct combinations of floral traits.

### Conclusions and management recommendations

Future research is needed to determine whether the effects of supplemental watering are consistent across additional native plant species, as indicated by the different responses of our two focal species. Assessing additional species will help determine if our results are consistent across taxonomic groups or co-flowering species, allowing for broader implementation of management recommendations that emerge from this work. Our interpretations were also limited by the length of our sampling period and the floral traits included in our study. Measuring additional floral characteristics, such as floral scent, pollen quality, or plant size, would help disentangle how supplemental watering affects different pollinator taxa. Sampling throughout the entire flowering season and over multiple years would help determine whether the effects of watering continue later in the season and if they are more pronounced over time. A more comprehensive understanding of the underlying mechanisms will help improve predictions of how supplemental watering influences plant-pollinator interactions under future climate conditions.

Our results provide practical insight into the management of urban green spaces in California and other regions where water conservation is a priority. Given that climate change is expected to increase the frequency and severity of drought (Diffenbaugh et al., 2015; Pachauri et al., 2014), developing water-efficient management strategies that simultaneously support pollinators is increasingly critical. Our findings suggest that supplemental water management can be tailored to plant phenology, allowing managers to prioritize supplemental water when most beneficial for pollinators. For spring-flowering plants, supplemental watering during spring may provide the greatest benefit to pollinator visitation, while summer-flowering plants may benefit from continued watering throughout summer. Although watering during spring only may provide some benefit to pollinator visitation of summer-flowering plants, extending watering into summer may maintain or increase this benefit as the season progresses. Matching supplemental water to flowering phenology may therefore allow managers to support pollinators while minimizing water use.

When extrapolated to other plant species, our findings imply that supplemental watering may preferentially benefit native bee communities. Native bees (bumblebees and Halictidae), rather than honeybees, were primarily driving the increase in pollinator visitation to spring-watered *P. californica*. In contrast, honeybee visitation was associated with variation in nectar sucrose concentrations instead of watering treatment, indicating that honeybees may rely on different and varied cues when selecting floral resources. Honeybees are a major competitor of native bees in urban areas, partially due to beekeeping (Egerer & Kowarik, 2020), and native bee diversity is threatened by habitat loss, urbanization, and climate change (Prendergast et al., 2022). Supplemental watering that promotes native plants in urban green spaces may therefore provide an additional conservation benefit to native bees.

Overall, our study revealed the ecological benefits of supplemental watering and potential management strategies to simultaneously minimize water use. By showing that the timing of water availability can influence plant-pollinator interactions, our study highlights opportunities to integrate pollinator conservation with water conservation, providing guidance for homeowners and stakeholders looking to build resilient urban habitats in an increasingly drought-prone climate.

## Author Contributions

Emily K. Meineke and Jaxi E. Cohen collaborated on study conception and design. Emily K. Meineke, Angelica M. Sauceda, Rachel M. Davis, Emily B. Griswold, and Lauren Azevedo-Schmidt collaborated on the design of the CALL Garden. All data collection (excluding phenology and corolla length) and analyses were performed by Jaxi E. Cohen. Phenology data collection was performed by Skye Austin. Emily K. Meineke facilitated the collection of corolla length measurements by undergraduate students as part of an undergraduate course. Rachel L. Vannette advised on nectar collection procedures. The first draft of the manuscript was written by Jaxi E. Cohen, and all authors commented on previous versions of the manuscript. All authors read and approved the final manuscript.

## Statement on inclusion

All authors are based at the University of California, Davis, in the region where the study was conducted. The research was carried out in collaboration with the UC Davis Arboretum and Public Gardens, whose staff are co-authors and contributed to study design, irrigation installation, plot construction, and plant care. Findings will be shared with the Arboretum’s public garden program to inform plantings and irrigation recommendations for visitors and local gardeners.

## Data availability statement

Data and code will be archived in Dryad upon acceptance.

## Acknowledgements

This material is based upon work supported by the U.S. National Science Foundation under Grant Number 2238310. Any opinions, findings, and conclusions or recommendations expressed in this material are those of the authors and do not necessarily reflect the views of the National Science Foundation.

