## Supplemental Information for "Supplemental water modulates pollinator visitation on resource-rich plant species"

**Supplementary Methods**

### S1. Irrigation system

Irrigation installation took place from October 2024 to February 2025. Plots were irrigated using surface drip irrigation with emitters spaced every 30.48 cm, each delivering 2.27 L $h^{-1}$. Surface irrigation lines were connected to a PVC manifold supplying water from four irrigation valves connected to a control box. Treatment-specific irrigation schedules were sent to the control box via Baseline software. To prevent water from flowing through all four irrigation valves simultaneously, the manifold was equipped with manual ball valves that were opened or closed depending on the treatment.

All irrigated plots received water for 3 h (05:00 to 08:00) every Monday and Friday, so treatments differed in the seasonal duration over which irrigation was applied, not in frequency. Weeding was done by hand throughout the experiment. Senesced flowers were removed after peak flowering of *P. californica* on July 23, 2025, and after 70% of flowers finished blooming on *H. puberulum* on October 29, 2025.

### S2. Plot construction and planting layout

Plots were constructed with 1.52 x 1.52 m wooden frames approximately 15.24 cm deep containing field soil (Yolo Slit Loam, Putah Creek Riparian Reserve) and a surface layer of mulch (Redi-Gro Corporation, Fir), consistent with how regional home gardeners often establish garden beds. Plots were spaced 45.72 cm apart. Each plot contained one individual of five native species: *Phacelia californica, Helenium puberulum*, *Artemisia douglasiana, Salvia mellifera,* and *Symphoricarpos albus* var. *laevigatus,* each centered within five quadrants of the plot. Species were planted in the same position within each plot. Only the first two species are analyzed here.

### S3. Sun and shade classification

During nectar sampling, we observed that the sun exposure of a plot was affecting the nectar production and sucrose concentration. Due to the foliage in the surrounding area, not all plots received equal shading in the morning. We assumed morning sun exposure was most relevant to our analysis, as it is when pollinator activity begins. To quantify variation in morning sun exposure across plots, we measured the surface temperatures of flowers using an infrared thermometer and compared plants in the shade versus sun. Because there was a significant difference in the surface temperatures ($\beta = 8.02$, $p < 0.005$), and there was no significant difference in surface temperature of plants of different water treatments ($P = 0.42$), we used a binary category of shade or sun to designate each plot's conditions in the morning. To assign designations to each plot, we recorded a time lapse on June 26, 2025, from before sunrise until all plots were in full sun, and observed the conditions of each plot at 9:00 AM, approximately when pollinator activity begins. Plots were categorized as "sun" or "shade" based on whether they were exposed to sun at this time. The same classification was applied to all sampling dates.

**S4. Nectar sampling**

Nectar sampling took place during May 2025. On each plant individual of *P. californica*, we selected five flowers for nectar production and sucrose concentration measurements. Chosen flowers were selected from flower clusters (cymes) previously covered with mesh bags to exclude pollinators. Cymes were covered before flower opening until flowering and collection. Collection occurred before 11:00 AM to reduce variability due to evaporation. We measured nectar production in each flower using 10 μl microcapillary tubes placed in contact with flower nectaries. Nectar concentration was measured using a handheld optical refractometer. Nectar traits were not measured for *H. puberulum* due to the small size of the flowers.

**S5. Corolla Length**

Data was collected using the following protocol developed as part of an undergraduate research course. Corolla length was measured in millimeters under a dissecting microscope. In each plot, three flowers were harvested from *P. californica*. Flowers were collected and measured over a series of 5 days within an 8-day collection period in May 2025, with corolla length recorded on the same day of collection. Under the dissecting microscope, utilizing forceps, we removed the sepals from all flowers and flattened the petals. Corolla length was not measured for *H. puberulum* due to small flower size.

**S6. Phenology**

Phenology sampling started on April 13, 2025, and continued approximately weekly until the end of the study (Table S1). The distribution of sampling time depended on the week. Early in the season, there were some weeks in which all plants could be surveyed in 1 day. Later in the season, we could not count flowers on all plants in 1 day, and thus surveys were spread out over 2-3 days.

For each survey, we counted all open flowering units in each plot. The flowering unit for each species was determined based on what was most intuitive for that species. For *P. californica*, we counted one inflorescence as one unit, and for *H. puberulum*, we counted one flower head as one unit . We only counted flowering units with at least one open flower. Reproductive units that included only unopened buds, wilted flowers or seed pods were not counted.

**S7. Soil moisture and water potential**

We monitored the soil moisture using METER TEROS 11 soil moisture sensors (METER Group, Inc. Pullman, WA, USA). Soil water potential was monitored with METER TEROS 21 soil water potential sensors. All sensors were installed at 30 cm soil depth. We installed one of each sensor type into two plots of each treatment type. Soil moisture and water potential measurements were automatically logged every 4 h by METER ZL6 advanced cloud data logger. Sensors and data loggers were installed in May 2025.

Sensor measurements were divided into two periods corresponding to the treatment watering schedule and sampling periods: before and after June 20, when irrigation for the spring-watered treatment ended. For each period and treatment, mean soil water content and water potential were calculated at the plot level and then summarized across plots as means ± standard deviation.

### Supplementary Tables

| Species | Survey dates (2025) |
|---|---|
| *P. californica* | April 13, April 18, April 21, April 28, May 4, May 11, May 25 |
| *H. puberulum* | April 21, April 28, May 4, June 25, July 1, July 7, July 21 |

**Table S1: Phenology survey dates for each species.**

| Period | Treatment | Water content ($m^3$ $m^{-3}$) | Water potential (kPa) |
|---|---|---|---|
| Before June 20 | watered | 0.22 ± 0.08 | -445 ± 682 |
| Before June 20 | unwatered | 0.14 ± 0.06 | -2021 ± 854 |
| After June 20 | spring-summer-watered | 0.18 ± 0.06 | -50.4 ± 30.2 |
| After June 20 | spring-watered | 0.21 ± 0.11 | -2631 ± 240 |
| After June 20 | unwatered | 0.13 ± 0.06 | -3297 ± 910 |

**Table S2: Soil water content and water potential (mean ± SD).** "Watered" before June 20, 2025 pools all irrigated treatments. Values are descriptive and were not statistically tested.

| Response Variable | Covariate Included | Plant Species | Scale | Predictor | Slope Estimate | SE | z/t value | *P* value |
|---|---|---|---|---|---|---|---|---|
| Total visitation | Nectar sucrose concentration | P.c. | Per Plant | Number of flowers | 0.281 | 0.032 | z = 8.77 | **<0.001*** ** |
| | | | | Nectar sucrose concentration | -0.063 | 0.040 | z = -1.57 | 0.116 |

| | | | | | | |
|---|---|---|---|---|---|---|
| | | Time of day | -0.044 | 0.035 | z = -1.24 | 0.216 |
| | | Sun/Shade | -0.230 | 0.093 | z = -2.46 | **0.014*** |
| | | Corolla length | 0.071 | 0.038 | z = 1.87 | 0.062 |
| | Per Flower | Number of flowers | -0.035 | 0.009 | t = -4.05 | **<0.001***** |
| | | Nectar sucrose concentration | -0.016 | 0.009 | t = -1.78 | 0.078 |
| | | Time of day | -0.008 | 0.009 | t = -0.84 | 0.405 |
| | | Sun/Shade | -0.037 | 0.024 | t = -1.53 | 0.130 |
| | | Corolla length | 0.013 | 0.010 | t = 1.36 | 0.176 |
| Halictidae | Per Plant | Number of flowers | 0.211 | 0.059 | z = 3.58 | **<0.001***** |
| | | Nectar sucrose concentration | -0.111 | 0.071 | z = -1.56 | 0.119 |
| | | Time of day | 0.074 | 0.054 | z = 1.38 | 0.168 |
| | | Sun/Shade | -0.386 | 0.216 | z = -1.79 | 0.074 |
| | | Corolla length | 0.102 | 0.089 | z = 1.16 | 0.247 |
| | Per Flower | Number of flowers | -0.021 | 0.006 | t = -3.21 | **0.002**** |
| | | Nectar sucrose concentration | -0.016 | 0.007 | t = -2.29 | **0.024*** |
| | | Time of day | 0.007 | 0.007 | t = 1.00 | 0.321 |
| | | Sun/Shade | -0.017 | 0.018 | t = -0.91 | 0.364 |
| | | Corolla length | 0.006 | 0.007 | t = 0.85 | 0.398 |

| | | | | | | |
|---|---|---|---|---|---|---|
| Bumblebee | Per Plant | Number of flowers | 0.313 | 0.051 | z = 6.20 | **<0.001*** ** |
| | | Nectar sucrose concentration | 0.047 | 0.062 | z = 0.77 | 0.445 |
| | | Time of day | -0.128 | 0.058 | z = -2.20 | **0.028*** |
| | | Sun/Shade | -0.076 | 0.154 | z = -0.49 | 0.622 |
| | | Corolla length | 0.023 | 0.061 | z = 0.37 | 0.712 |
| | Per Flower | Number of flowers | -0.006 | 0.004 | t = -1.76 | 0.081 |
| | | Nectar sucrose concentration | 0.005 | 0.004 | t = 1.19 | 0.239 |
| | | Time of day | -0.008 | 0.004 | t = -2.10 | **0.038*** |
| | | Sun/Shade | -0.013 | 0.010 | t = -1.24 | 0.217 |
| | | Corolla length | -0.001 | 0.004 | t = -0.25 | 0.804 |
| Honeybee | Per Plant | Number of flowers | 0.228 | 0.104 | z = 2.19 | **0.028*** |
| | | Nectar sucrose concentration | -0.347 | 0.153 | z = -2.27 | **0.023*** |
| | | Time of day | -0.058 | 0.103 | z = -0.57 | 0.572 |
| | | Sun/Shade | -0.203 | 0.361 | z = -0.56 | 0.575 |
| | | Corolla length | 0.082 | 0.153 | z = 0.54 | 0.591 |
| | Per Flower | Number of flowers | -0.008 | 0.003 | t = -2.76 | **0.007**** |

| | | | | | | | | |
|---|---|---|---|---|---|---|---|---|
| | | | | Nectar sucrose concentration | -0.007 | 0.003 | t = -2.08 | **0.040*** |
| | | | | Time of day | -0.002 | 0.003 | t = -0.74 | 0.460 |
| | | | | Sun/Shade | -0.006 | 0.009 | t = -0.76 | 0.448 |
| | | | | Corolla length | 0.005 | 0.003 | t = 1.55 | 0.124 |
| Taxonomic richness | | | Per Plant | Number of flowers | 0.166 | 0.056 | z = 2.95 | **0.003**** |
| | | | | Nectar sucrose concentration | -0.048 | 0.067 | z = -0.72 | 0.472 |
| | | | | Time of day | -0.086 | 0.061 | z = -1.39 | 0.164 |
| | | | | Sun/Shade | -0.051 | 0.164 | z= -0.31 | 0.758 |
| | | | | Corolla length | 0.049 | 0.065 | z= 0.75 | 0.451 |
| Total visitation | Nectar volume | P.c. | Per Plant | Number of flowers | 0.281 | 0.032 | z = 8.77 | **<0.001***** |
| | | | | Nectar volume | -0.014 | 0.037 | z = -0.37 | 0.710 |
| | | | | Time of day | -0.057 | 0.035 | z = -1.64 | 0.102 |
| | | | | Sun/Shade | -0.269 | 0.092 | z = -2.93 | **0.003**** |
| | | | | Corolla length | 0.076 | 0.038 | z = 2.01 | **0.045*** |
| | | | Per Flower | Number of flowers | -0.035 | 0.009 | t = -3.97 | **<0.001***** |
| | | | | Nectar volume | -0.006 | 0.009 | t = -0.69 | 0.490 |
| | | | | Time of day | -0.012 | 0.009 | t = -1.35 | 0.179 |
| | | | | Sun/Shade | -0.049 | 0.024 | t = -2.04 | **0.043*** |
| | | | | Corolla length | 0.015 | 0.010 | t = 1.55 | 0.123 |

| | | | | | | |
|---|---|---|---|---|---|---|
| Halictidae | Per Plant | Number of flowers | 0.195 | 0.06 | z = 3.28 | **0.001**** |
| | | Nectar volume | 0.007 | 0.067 | z = 0.10 | 0.920 |
| | | Time of day | 0.054 | 0.052 | z = 1.04 | 0.299 |
| | | Sun/Shade | -0.457 | 0.217 | z = -2.11 | **0.035*** |
| | | Corolla length | 0.113 | 0.090 | z = 1.25 | 0.212 |
| | Per Flower | Number of flowers | -0.021 | 0.007 | t = -3.14 | **0.002**** |
| | | Nectar volume | -0.002 | 0.007 | t = -0.27 | 0.789 |
| | | Time of day | 0.003 | 0.007 | t = 0.41 | 0.684 |
| | | Sun/Shade | -0.027 | 0.018 | t = -1.49 | 0.139 |
| | | Corolla length | 0.008 | 0.007 | t = 1.04 | 0.299 |
| Bumblebee | Per Plant | Number of flowers | 0.313 | 0.051 | z = 6.19 | **<0.001***** |
| | | Nectar volume | -0.007 | 0.060 | z = -0.11 | 0.912 |
| | | Time of day | -0.118 | 0.056 | z = -2.09 | **0.037*** |
| | | Sun/Shade | -0.052 | 0.151 | z = -0.35 | 0.730 |
| | | Corolla length | 0.021 | 0.061 | z = 0.34 | 0.734 |
| | Per Flower | Number of flowers | -0.006 | 0.004 | t = -1.72 | 0.089 |
| | | Nectar volume | -0.002 | 0.004 | t = -0.44 | 0.659 |
| | | Time of day | -0.007 | 0.004 | t = -1.87 | 0.064 |
| | | Sun/Shade | -0.01 | 0.010 | t = -1.02 | 0.312 |
| | | Corolla length | -0.001 | 0.004 | t = -0.31 | 0.756 |

| | | | | | | | |
|---|---|---|---|---|---|---|---|
| Honeybee | | Per Plant | Number of flowers | 0.193 | 0.106 | z = 1.82 | 0.069 |
| | | | Nectar volume | 0.023 | 0.129 | z = 0.18 | 0.855 |
| | | | Time of day | -0.097 | 0.102 | z = -0.96 | 0.338 |
| | | | Sun/Shade | -0.386 | 0.370 | z = -1.04 | 0.297 |
| | | | Corolla length | 0.104 | 0.157 | z = 0.66 | 0.510 |
| | | Per Flower | Number of flowers | -0.008 | 0.003 | t = -2.70 | **0.008**** |
| | | | Nectar volume | -0.001 | 0.003 | t = -0.40 | 0.690 |
| | | | Time of day | -0.004 | 0.003 | t = -1.31 | 0.194 |
| | | | Sun/Shade | -0.011 | 0.008 | t = -1.30 | 0.195 |
| | | | Corolla length | 0.006 | 0.003 | t = 1.73 | 0.086 |
| Taxonomic richness | | Per Plant | Number of flowers | 0.167 | 0.057 | z = 2.95 | **0.003**** |
| | | | Nectar volume | -0.014 | 0.062 | z = -0.23 | 0.819 |
| | | | Time of day | -0.097 | 0.059 | z = -1.61 | 0.107 |
| | | | Sun/Shade | -0.081 | 0.162 | z = -0.50 | 0.615 |
| | | | Corolla length | 0.054 | 0.065 | z = 0.82 | 0.451 |
| Total visitation NA | H.p. | Per Plant | Number of flowers | 141 | 18.4 | t = 7.67 | **<0.001***** |
| | | | Time of day | -12.6 | 23.5 | t = -0.54 | 0.593 |
| | | | Sun/Shade | 53.2 | 58.4 | t = 0.91 | 0.364 |
| | | Per Flower | Number of flowers | 0.388 | 0.068 | z = 5.70 | **<0.001***** |

| | | | | | |
|---|---|---|---|---|---|
| | Time of day | 0.125 | 0.097 | z = 1.29 | 0.198 |
| | Sun/Shade | -0.164 | 0.279 | z = -0.59 | 0.556 |

**Table S3: Mechanistic model results.** *P. californica* per plant and *H. puberulum* perflower visitation, and *P. californica* richness: Poisson GLMMs (log-scale estimates, *z*). *P. californica* per flower and *H. puberulum* per-plant visitation: Gaussian LMMs (estimates in visits per plant per 3 min, *t*). * $p < 0.05$, ** $p < 0.01$, and *** $p < 0.001$. *P.c., Phacelia californica; H.p., Helenium puberulum.*

| **Analysis** | **Response variable** | **Species** | **Fixed effects** | **Random effect** | **Model** | **Distribution** | **n** |
|---|---|---|---|---|---|---|---|
| Supplemental water effects on visitation | Total visitation per plant | *P. c.* | treatment x date; sun exposure; time | plot | GLMM | Poisson | 119 |
| Supplemental water effects on visitation | Total visitation per flower | *P. c.* | treatment x date; sun exposure; time | plot | LMM | Gaussian | 119 |
| Supplemental water effects on visitation | Bumblebee visitation per plant | *P. c.* | treatment x date; sun exposure; time | plot | GLMM | Poisson | 119 |
| Supplemental water effects on visitation | Bumblebee visitation per flower | *P. c.* | treatment x date; sun exposure; time | plot | LMM | Gaussian | 119 |
| Supplemental water effects on visitation | Honeybee visitation per plant | *P. c.* | treatment x date; sun exposure; time | plot | GLMM | Poisson | 119 |
| Supplemental water effects on visitation | Honeybee visitation per flower | *P. c.* | treatment x date; sun exposure; time | plot | LMM | Gaussian | 119 |
| Supplemental water effects on visitation | Halictidae visitation per plant | *P. c.* | treatment x date; sun exposure; time | plot | GLMM | Poisson | 119 |

| | | | | | | | |
|---|---|---|---|---|---|---|---|
| Supplemental water effects on visitation | Halictidae visitation per flower | *P. c.* | treatment x date; sun exposure; time | plot | LMM | Gaussian | 119 |
| Supplemental water effects on category richness | Category richness per plant | *P. c.* | treatment x date; sun exposure; time | plot | GLMM | Poisson | 119 |
| Mechanistic effects on visitation | Total visitation per plant | *P. c.* | number of flowers; mean corolla length; sun exposure, time of day, mean nectar volume | plot | GLMM | Poisson | 119 |
| Mechanistic effects on visitation | Total visitation per plant | *P. c.* | number of flowers; mean corolla length; sun exposure, time of day, nectar sucrose concentration | plot | GLMM | Poisson | 119 |
| Mechanistic effects on visitation | Total visitation per flower | *P. c.* | number of flowers; mean corolla length; sun exposure, time of day, mean nectar volume | plot | LMM | Gaussian | 119 |
| Mechanistic effects on visitation | Total visitation per flower | *P. c.* | number of flowers; mean corolla length; sun exposure, time of day, nectar sucrose concentration | plot | LMM | Gaussian | 119 |

| | | | | | | | |
|---|---|---|---|---|---|---|---|
| Mechanistic effects on visitation | Bumblebee visitation per plant | *P. c.* | number of flowers; mean corolla length; sun exposure, time of day, mean nectar volume | plot | GLMM | Poisson | 119 |
| Mechanistic effects on visitation | Bumblebee visitation per plant | *P. c.* | number of flowers; mean corolla length; sun exposure, time of day, nectar sucrose concentration | plot | GLMM | Poisson | 119 |
| Mechanistic effects on visitation | Bumblebee visitation per flower | *P. c.* | number of flowers; mean corolla length; sun exposure, time of day, mean nectar volume | plot | LMM | Gaussian | 119 |
| Mechanistic effects on visitation | Bumblebee visitation per flower | *P. c.* | number of flowers; mean corolla length; sun exposure, time of day, nectar sucrose concentration | plot | LMM | Gaussian | 119 |
| Mechanistic effects on visitation | Honeybee visitation per plant | *P. c.* | number of flowers; mean corolla length; sun exposure, time of day, mean nectar volume | plot | GLMM | Poisson | 119 |

| | | | | | | | |
|---|---|---|---|---|---|---|---|
| Mechanistic effects on visitation | Honeybee visitation per plant | *P. c.* | number of flowers; mean corolla length; sun exposure, time of day, nectar sucrose concentration | plot | GLMM | Poisson | 119 |
| Mechanistic effects on visitation | Honeybee visitation per flower | *P. c.* | number of flowers; mean corolla length; sun exposure, time of day, mean nectar volume | plot | LMM | Gaussian | 119 |
| Mechanistic effects on visitation | Honeybee visitation per flower | *P. c.* | number of flowers; mean corolla length; sun exposure, time of day, nectar sucrose concentration | plot | LMM | Gaussian | 119 |
| Mechanistic effects on visitation | Halictidae visitation per plant | *P. c.* | number of flowers; mean corolla length; sun exposure, time of day, mean nectar volume | plot | GLMM | Poisson | 119 |
| Mechanistic effects on visitation | Halictidae visitation per plant | *P. c.* | number of flowers; mean corolla length; sun exposure, time of day, nectar sucrose concentration | plot | GLMM | Poisson | 119 |

| | | | | | | | |
|---|---|---|---|---|---|---|---|
| Mechanistic effects on visitation | Halictidae visitation per flower | *P. c.* | number of flowers; mean corolla length; sun exposure, time of day, mean nectar volume | plot | LMM | Gaussian | 119 |
| Mechanistic effects on visitation | Halictidae visitaton per flower | *P. c.* | number of flowers; mean corolla length; sun exposure, time of day, nectar sucrose concentration | plot | LMM | Gaussian | 119 |
| Mechanistic effects on visitation | Category richness per plant | *P. c.* | number of flowers; mean corolla length; sun exposure, time of day, mean nectar volume | plot | GLMM | Poisson | 119 |
| Mechanistic effects on visitation | Category richness per plant | *P. c.* | number of flowers; mean corolla length; sun exposure, time of day, nectar sucrose concentration | plot | GLMM | Poisson | 119 |
| Supplemental water effects on visitation | Total visitation per plant | *H. p.* | treatment x date; sun exposure; time of day | plot | LMM | Gaussian | 160 |
| Supplemental water effects on visitation | Total visitation per flower | *H. p.* | treatment x date; sun exposure; time of day | plot | GLMM | Poisson | 160 |

| | | | | | | | |
|---|---|---|---|---|---|---|---|
| Mechanistic effects on visitation | Total visitation per plant | *H. p.* | number of flowers; sun exposure, time of day | plot | LMM | Gaussian | 160 |
| Mechanistic effects on visitation | Total visitation per flower | *H. p.* | number of flowers; sun exposure, time of day | plot | GLMM | Poisson | 160 |
| Sun exposure effects on surface temperatere | Surface temperature | *NA* | sun exposure, treatment | plot; date | LMM | Gaussian | 90 |
| Supplemental water effects on flower traits | Number of flowers | *P. c.* | treatment x date; sun exposure | plot | LMM | Gaussian | 280 |
| Supplemental water effects on flower traits | Number of flowers | *H. p.* | treatment x date; sun exposure | plot | LMM | Gaussian | 201 |
| Supplemental water effects on flower traits | Nectar sucrose concentration | *P. c.* | treatment x date; sun exposure; mean corolla length; nectar volume | plot | LMM | Gaussian | 80 |
| Supplemental water effects on flower traits | Nectar volume | *P. c.* | treatment x date; sun exposure; mean corolla length; nectar sucrose concentration | plot | LMM | Gaussian | 80 |

**Table S4: List of models.** Response variables are total visitation per plant or per flower, flower traits, or flower surface temperature. *n* is the number of observations in each model. *P.c., Phacelia californica; H.p., Helenium puberulum.*
